# Dynamic Kinetochore Size Regulation Promotes Microtubule Capture and Chromosome Biorientation in Mitosis

**DOI:** 10.1101/279398

**Authors:** Carlos Sacristan, Misbha Ahmad, Jenny Keller, Job Fermie, Vincent Groenewold, Eelco Tromer, Alexander Fish, Roberto Melero, José María Carazo, Judith Klumperman, Andrea Musacchio, Anastassis Perrakis, Geert JPL Kops

## Abstract

Faithful chromosome segregation depends on the ability of sister kinetochores to attach to spindle microtubules. An outer layer of the kinetochore known as the fibrous corona transiently expands in early mitosis and disassembles upon microtubule capture. Neither the functional importance nor the mechanistic basis for this are known. Here we show that the dynein adaptor Spindly and the RZZ kinetochore complex drive fibrous corona formation in a dynein-independent manner. C-terminal farnesylation and MPS1 kinase activity cause conformational changes of Spindly that promote oligomerization of RZZ:Spindly complexes into a corona-like meshwork in cells and *in vitro*. Concurrent with corona expansion, Spindly potentiates corona shedding by recruiting dynein via three conserved short linear motifs. Expanded, non-sheddable fibrous coronas engage in extensive, long-lived lateral microtubule interactions that persist to metaphase and result in fused sister kinetochores, formation of merotelic attachments and chromosome segregation errors in anaphase. Thus, dynamic kinetochore size regulation in mitosis is coordinated by a single, Spindly-based mechanism that promotes initial microtubule capture and subsequent correct maturation of attachments.

## INTRODUCTION

The kinetochore is a complex protein structure that physically anchors the chromosomes to spindle microtubules to ensure chromosome segregation during cell division ^1,2^. Defects in kinetochore function cause chromosome segregation errors that lead to aneuploid cellular progeny. Aneuploidy is a major cause of embryonic maldevelopment and a prominent genomic abnormality in human cancers ^3–9^.

To guarantee the fidelity of chromosome segregation in mitosis, chromosomes must capture microtubules and achieve stable bioriented attachments before anaphase, meaning they are bound to microtubules from opposite spindle poles. Microtubule capture and biorientation are complex processes that require an intricate kinetochore machinery, involving dozens of proteins that dynamically adapt to the microtubule attachment status of the kinetochore ^1,2,10–12^. The core of the kinetochore in animals and fungi is made of the CCAN and the KMN network. The CCAN associates with centromeric chromatin and connects it to the microtubule-binding site that resides within the HEC1/NDC80 and NUF2 members of the KMN network ^2^. HEC1/NDC80 is a key target in the biorientation pathway, as its phosphorylation by Aurora B kinase signals non-bioriented attachments that are then corrected accordingly ^13^.

The core kinetochore module is supplemented with proteins whose quantity at kinetochores changes during mitotic progression. These include proteins of the spindle assembly checkpoint (SAC), which bind predominantly to the scaffold KNL1 when HEC1/NDC80 is not bound to microtubules ^14^. Other transient kinetochore proteins, more distal to the centromeric chromatin, are dynein recruiters (ROD-ZW10-Zwilch (RZZ) and Spindly), dynein regulators (CENP-F/Nude1/Nde1/CLIP-170), the kinesins CENP-E and Kif2b, and modifiers of microtubule dynamics such as CLASP1 and CLASP2 ^15,16^. These proteins are involved in initial microtubule capture, chromosome transport, and microtubule-based force generation that enables, amongst others, anaphase.

The mammalian kinetochore exhibits great morphological plasticity during mitosis. While the core kinetochore module expands little if any (with the exception of CENP-C in *Xenopus* extracts), the more centromere-distal modules expand to large, crescent shapes in early prometaphase and collapse into compact spherical structures in metaphase ^17–27^. Electron micrographs of kinetochores in their most expanded form show a halo of low-density material referred to as the fibrous corona, which is absent from kinetochores that are bound by microtubules ^28–33^. Neither the molecular nature of the fibrous corona nor its functional relevance are known. Recent computational modeling suggested that the fibrous corona promotes spindle assembly, and that correct rotation of the sister kinetochores followed by their compaction then reduces the risk of erroneous attachments ^17,34^. Experimental approaches to manipulate the fibrous corona in order to test this and other hypotheses are lacking ^34^.

The dynamic behaviour of the fibrous corona correlates with that of the proteins that display early mitotic expansion and subsequent compaction. It is therefore likely that one or more of them can form the corona meshwork, and that regulation of its expansion/compaction will impinge on these components. A candidate is the RZZ complex. RZZ shows classic expansion/compaction behavior, is required for recruitment of several outer-kinetochore components, and has a molecular architecture resembling that of membrane coating proteins that can form polymeric states ^25,35–44^. We here set out to identify the nature of the fibrous corona,the molecular mechanisms of its ability to expand and compact, and its functional relevance for chromosome segregation. We show that the Spindly protein is at the heart of fibrous corona biology: It forms the fibrous corona by triggering polymerization of RZZ-Spindly complexes, and subsequently sets the stage for later corona compaction by recruiting the dynein motor complex. In this way, Spindly couples the mechanisms of physical expansion of kinetochores with subsequent compaction and we show this enables efficient capture and stabilization of correct microtubule attachments.

## RESULTS

### Spindly recruits dynein to compact kinetochores upon microtubule attachment

In agreement with previous observations ^17,18,22^, kinetochores expand in nocodazole-treated cells shortly after NEB, as evidenced by the formation of ZW10-positive crescents (Figure 1a, examples 1 to 3). The crescents grew during mitosis and merged with those of the sister kinetochore to form large, ring-shaped structures (Figure 1a, example 4). Rings were absent from cells in which a spindle was allowed to form, and the attached kinetochore on a mono-oriented chromosome was more compact than its unattached sister kinetochore (Figure 1a, example 5). To examine the function of this expansion and subsequent compaction by microtubules, we wished to understand its molecular underpinnings. In *Xenopus* egg extracts, kinetochore size is regulated by mitotic phosphorylation ^24^. Phosphatases were, however, unable to reverse kinetochore expansion in human cells, as small molecule inhibitors of Aurora B, PLK1 or MPS1, when added after expansion, did not cause compaction of kinetochores (Figure S1).

**Figure 1:**
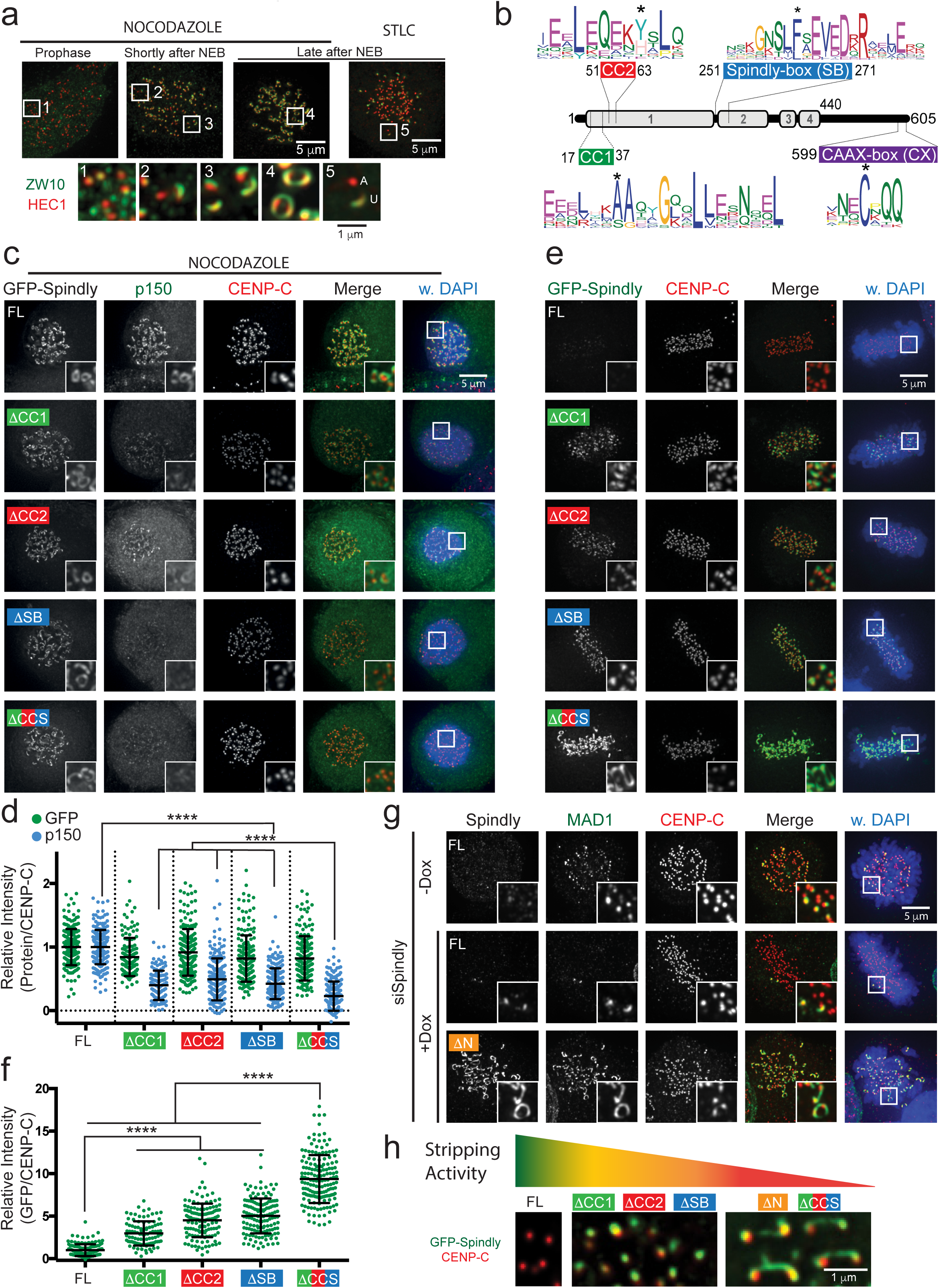
Spindly recruits dynein to compact kinetochores upon microtubule attachment. (a) Immunofluorescence images of ZW10 and HEC1 in HeLa cells treated with nocodazole or STLC. A, Attached; U, Unattached. **(b)** Overview of the secondary structure of human Spindly with predicted coiled-coils (grey bars) and disordered regions, and with sequence logos of four conserved motifs (supplementary sequences 1). **(c,d)** Representative images (c) and quantification (d) of nocodazole-treated HeLa cells transfected with siRNAs to Spindly and expressing the indicated GFP-Spindly variants, and immuno-stained for the indicated antigens. The graph in D shows mean kinetochore intensity (±s.d.) normalized to the values of Spindly^FL^ from at least four independent experiments. Each dot represents one cell and at least 20 cells were quantified per experiment and condition. Asterisks indicate significance (analysis of variance with Tukey’s multiple comparison test). ****P < 0.001. **(e,f)** Representative images (e) and quantification (f) of metaphase HeLa cells transfected with siRNA to Spindly and expressing the indicated GFP-Spindly variants, and immuno-stained for the indicated antigens. The graph shows the mean fold change in kinetochore intensity (±s.d.) normalized to the values of Spindly^FL^ from five independent experiments. Each dot represents one cell and at least 20 cells were quantified per experiment and condition. Asterisks indicate significance (analysis of variance with Tukey’s multiple comparison test). ****P < 0.001. **(g)** Immunofluorescence images of HeLa cells transfected with siRNA to Spindly and treated or not with doxycycline (Dox) to induce the expression of GFP-Spindly variants, and immuno-stained for the indicated antigens. **(h)** Representative images of the morphology of Spindly and CENP-C in metaphase kinetochores expressing the indicated GFP-Spindly variants. Stripping activity is based on ability of Spindly variants to recruit dynein/dynactin.

Dynein drives poleward transport of kinetochore proteins ^25,45–50^. To investigate if dynein is required for compaction of expanded kinetochores, we examined Spindly, a predicted coiled-coil protein that is the dynein/dynactin adaptor at kinetochores ^26,35,43,51–53^. Spindly localization to kinetochores requires a conserved C-terminal CAAX box (Figure 1b), whose farnesylation promotes binding to the RZZ complex ^35,43,51,54,55^. Spindly in turn recruits dynein/dynactin through two motifs: the Spindly box (SB), and the recently identified CC1 box (Figure 1b and S2a) ^35,43,51,52^. Our ConFeaX pipeline, which identifies conserved motifs in orthologous protein families ^56,57^, additionally identified a Qxx[HY] motif close to the CC1 box, which we here named the CC2 box (Figure 1b). CC2-like boxes can be found in other dynein/dynactin adaptors (BICD1/2, BICDR1/2, TRAK1/2, and HAP1, Figure S2a), suggesting they are involved in dynein/dynactin binding. Indeed, recruitment of the dynactin subunit p150^glued^ to kinetochores was compromised in cells expressing GFP-Spindly mutated in either of the three motifs [A24V (Spindly^ΔCC1^), Y60A (Spindly^ΔCC2^), F258A (Spindly^ΔSB^)], and, importantly, was nearly abolished when all three motifs were mutated (Spindly^ΔCCS^) (Figure 1c,d). Dynein recruitment by Spindly results in stripping of Spindly from kinetochores upon microtubule attachment ^26,43,51–53,58,59^. In accordance with this, the low kinetochore dynactin levels in cells expressing Spindly motif mutants caused those mutants to be retained on metaphase kinetochores (Figure 1e,f). Strikingly, Spindly^ΔCCS^, which was the most compromised in recruiting dynein (Figure 1e, f), frequently appeared as a large structure bridging the two sister kinetochores (Figure 1e). This resembled fully expanded kinetochores of nocodazole-treated cells (compare example 4 in Figure 1a with inset of Spindly^ΔCCS^ in Figure 1e). A similar phenotype was observed by deleting the N-terminal 65 amino acids of Spindly (Spindly^ΔN^) that contain the CC1 and CC2 boxes (Figure 1g). These findings suggest that Spindly, via the recruitment of dynein, is essential to compact previously expanded kinetochores (Figure 1h).

### Kinetochores expand by forming a structurally stable fibrous corona

Having established the mechanism of kinetochore compaction, we next examined the molecular basis for the kinetochore expansion. Quantitative immunofluorescence imaging of kinetochores in nocodazole-treated cells showed that the RZZ complex, Spindly and MAD1 occupied the largest volumes (often showing partial or complete encirclement of the sister kinetochores), followed by the KMN network and the CCAN (Figure 2a,b). This is in agreement with studies reporting that different kinetochore modules expand to different extents and inversely to their proximity to the chromosome’s axis ^17,18,24^. RZZ, Spindly and MAD1 are thought to be part of the fibrous corona, which in electron micrographs appears as a thick electron-dense structure that assembles on the outer kinetochore in early mitosis or upon addition of microtubule-deplymerizing drugs ^17,19,21,22,25,28,30,33,36,42,55^. We thus deemed it likely that kinetochore expansion observed in immunofluorescence images represents corona assembly. RZZ, Spindly and MAD1 may therefore constitute the core fibrous corona meshwork and/or regulate its formation. This was supported by the observation that brief (20’) CDK1 inhibition caused the detachment from the inner parts of the kinetochore of intact, crescent-shaped rods or extended filaments close to the kinetochores that contained Spindly, ZW10, MAD1 and CENP-E (Figure 2c). These data support the hypothesis that the expanded kinetochore module observed in the absence of microtubules is the fibrous corona and that it is a stable structure with a specific architecture, composition and regulation, distinct from other outer kinetochore modules such as the KMN network (Figure S3a).

**Figure 2:**
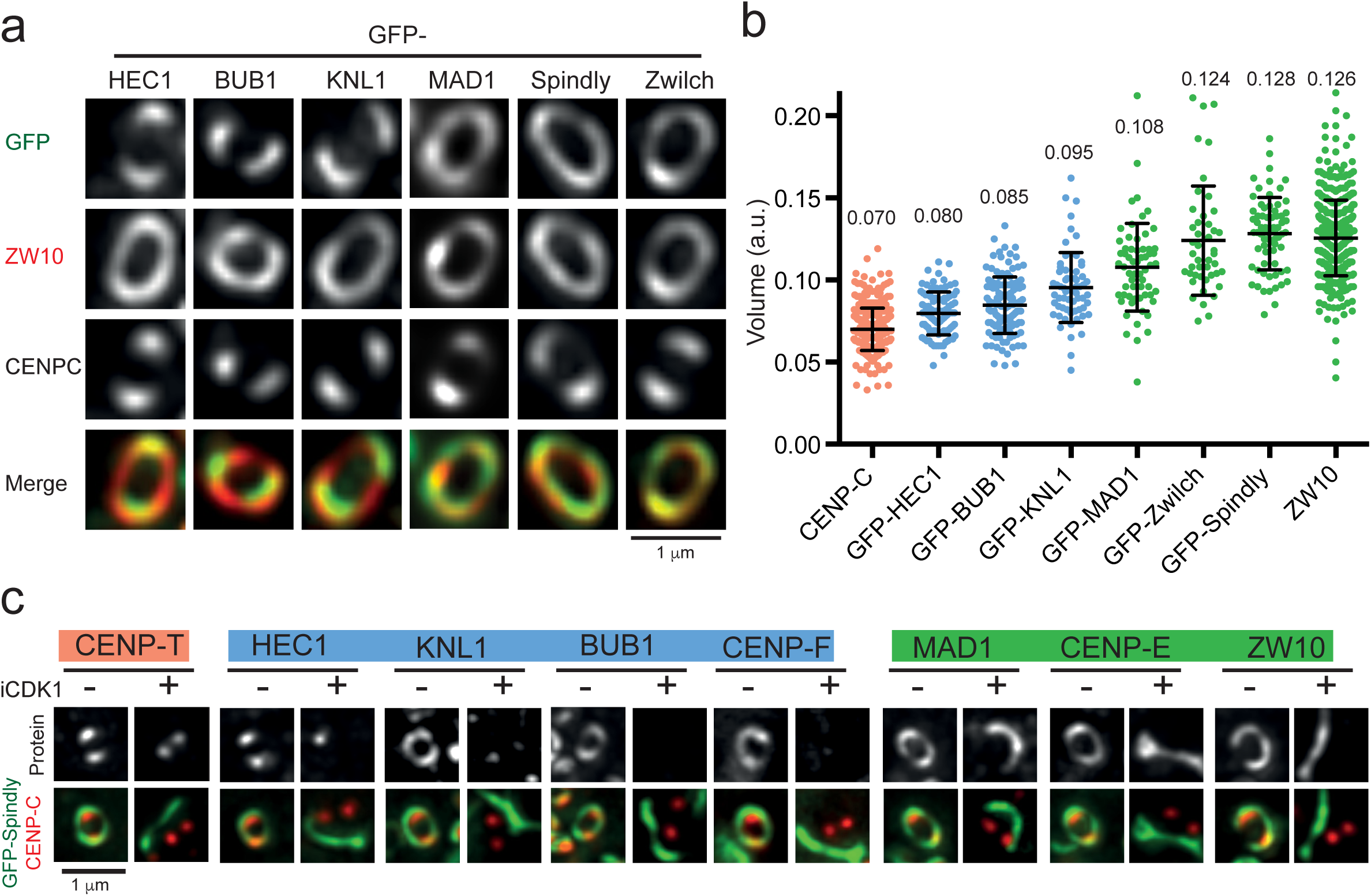
Kinetochores expand by forming a structurally stable fibrous corona. (a,b) Representative images (a) and volume quantification (b) of the indicated GFP-tagged proteins in expanded kinetochores of HeLa cells entering mitosis in the presence of nocodazole. Maximum expanded kinetochores were selected based on ZW10 staining. The graph shows the mean kinetochore volume (±s.d.) of at least three independent experiments. Each dot represents one kinetochore and at least 10 kinetochores per experiment were quantified. **(c)** Images of kinetochores of HeLa cells entering mitosis in the presence of nocodazole and subsequently treated with RO-3306 for 20 minutes, and immunostained with the indicated antibodies. Color codes in B and C correspond to behavior of protein after brief CDK1 inhibition.

### Spindly and RZZ are essential for corona formation

Immunofluorescence imaging and electron microscopy showed absence of kinetochore expansion and absence of a fibrous corona, respectively, in ZW10 RNAi cells (Figure 3a,c,e,g and Figure S3b-d), further establishing that kinetochore expansion equates to formation of a fibrous corona. Interestingly, ZW10 RNAi also reduced the volumes occupied by the KMN network member HEC1 and the CCAN member CENP-C (Figure 3a, c). RZZ is thus essential for fibrous corona formation, and controls the (limited) expansion of inner- and outer- kinetochore regions.

**Figure 3:**
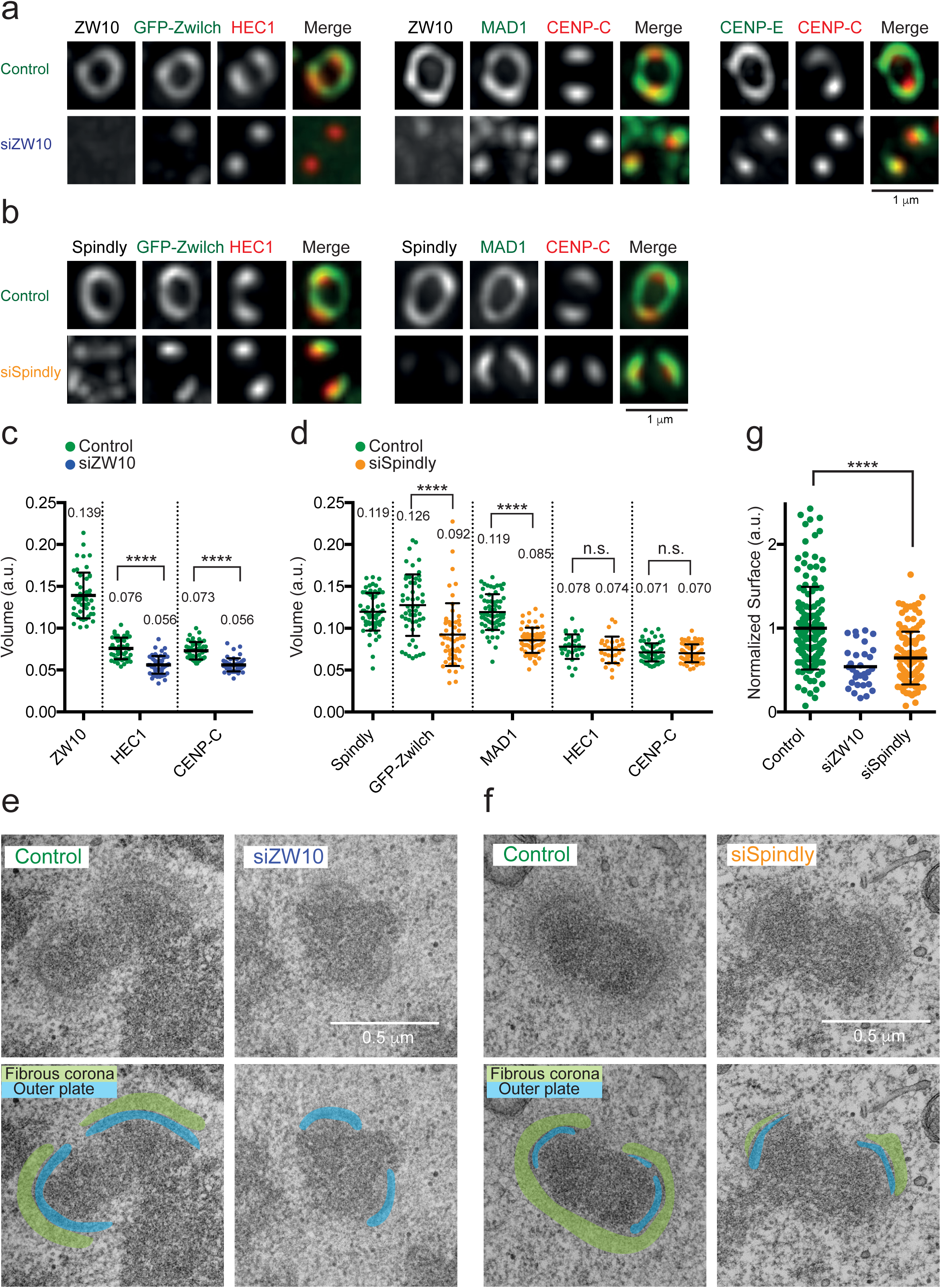
Spindly and RZZ are essential for corona formation. (a-d) Representative images (a, b) and volume quantification (c, d) based on the indicated antigens of expanded kinetochores of nocodazole-treated HeLa cells transfected with siRNA to ZW10 (a,c) or Spindly (b,d). The graph shows the mean kinetochore volume (±s.d.) of at least three independent experiments. Each dot represents one kinetochore and at least 10 kinetochores per experiment were quantified. Asterisks indicate significance (analysis of variance with Tukey’s multiple comparison test). ****P < 0.001; n.s., not significant. **(e-g)** Electron micrographs (e,f) and surface quantification (g) of the fibrous corona of nocodazole-treated cells depleted of ZW10 (e) or Spindly (f). The fibrous corona is highlighted in green and the outer plate in blue. The graph shows the mean surface of the fibrous corona (±s.d.) normalized to the value of the control from two (control and siSpindly) or one (siZW10) experiment. At least 50 kinetochores per condition and experiment were imaged. Quantification of the protein depletion efficiency are shown in Figure S3c,d. Each dot represents a single kinetochore. Asterisks indicate significance (Student’s t-test, unpaired). ****P < 0.001. Control is siGAPDH.

We next turned our attention to Spindly, and reasoned that if dynein recruitment were its predominant contribution to kinetochore size dynamics, Spindly depletion should phenocopy severe Spindly motif mutants, i.e. persistently expanded kinetochores due to an inability of microtubules to cause shedding of the corona (Figure 1g). In contrast, however, volume measurements and electron microscopy revealed that cells depleted of Spindly were unable to expand kinetochores and had severely compromised fibrous coronas (Figure 3b,d,f,g and Figure S3b-d). Spindly is therefore essential not only for corona disassembly through dynein binding, but also for corona assembly.

### Spindly stimulates RZZ-Spindly polymerization *in vitro* and *in vivo*

RZZ is necessary for the recruitment of several fibrous corona components ^22,26,37,38,40,53,58,60^, and its structure resembles that of coat scaffolds, which can self-assemble into polymeric states ^35,44^. We therefore asked if RZZ could assemble ectopically into polymeric structures, which may be surmised to promote the expansion of the corona. Purified, recombinant RZZ, which assembles with 2:2:2 (ROD:Zwilch:ZW10) stoichiometry^35^, did not oligomerize, as assessed by direct visualization of mCherry-ROD by fluorescence microscopy (Figure 4a). Addition of purified farnesylated Spindly (Spindly^FAR^), however, caused spontaneous oligomerization into filamentous structures at 30°C (Figure 4c, Movie S1). *In vitro* filament formation of RZZ-Spindly (RZZS) complexes could be prevented by addition of detergent, suggesting that hydrophobic interactions underlie the assembly reaction (Figure 4a). Notably, RZZS oligomerization in the presence of GFP-Spindly-coated agarose beads resulted in stable association of filaments with the beads (Figure 4b), in a configuration reminiscent of the fibrous corona at kinetochores.

**Figure 4:**
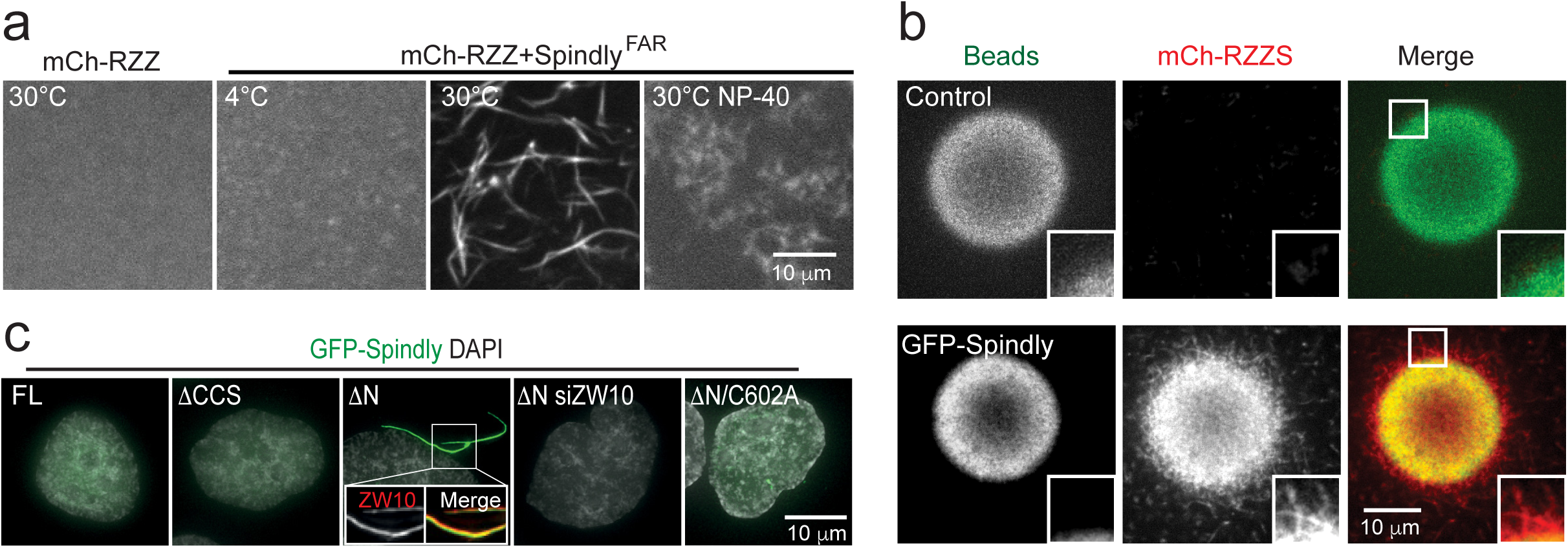
Spindly stimulates RZZ-Spindly polymerization *in vitro* and *in vivo*. (a) Fluorescence microscopy of purified mCherry-RZZ alone or in the presence of recombinant farnesylated Spindly (Spindly^FAR^) and incubated as indicated. See alsoMovie S1. **(b)** GFP-Spindly immobilized on beads and incubated in the presence of mCherry-RZZ and farnesylated Spindly (RZZS). Control is empty beads. The intensity level of the green channel in the control were enhanced to allow the comparison with GFP-Spindly. **(c)** Immunofluorescence images of interphase HeLa cells expressing the indicated versions of GFP-Spindly and treated and stained as indicated. See also Movie S2.

Expression of Spindly^FL^ (full length) in interphase cells had no discernable effect on RZZ complexes, but expression of Spindly^ΔN^ caused spontaneous formation of cytoplasmic filaments containing ZW10, Zwilch and ROD (Figure 4c, S4a and Movie S2). These filaments strongly resembled the ones formed *in vitro* by purified components and were strikingly similar to the detached coronas apparent after brief CDK1 inhibition (see Figure 2c). Cytoplasmic filament formation was abolished upon ZW10 RNAi or mutation of the Spindly CAAX box (C602A), showing it relied on RZZ and RZZ’s interaction with farnesylated Spindly (Figure 4c). Spindly^ΔCCS^, which like Spindly^ΔN^ is unable to bind dynein/dynactin, did not induce filaments (Figure 4c), showing that the ability of Spindly^ΔN^ to cause RZZS oligomerization is unrelated to dynein/dynactin binding.

### Spindly adopts a 3D configuration that impairs its ability to promote RZZS oligomerization

Collectively, these results indicate that Spindly can promote the formation of a stable RZZS meshwork, and that this process is greatly enhanced by deletion of the N-terminal 65 amino acids of Spindly. To examine how removal of the N-terminal 65 amino acids of Spindly promotes cytoplasmic RZZ filament formation, we next wished to obtain a better understanding of Spindly’s structure and regulation. Residues 1-440 of Spindly are predicted to form a coiled-coil structure while the C-terminal region is likely disordered (Figure 1b) ^35,43,54^. Small Angle X-ray Scattering (SAXS) analysis of a recombinant Spindly fragment encompassing 1-440 showed intramolecular dimensions (D_max_, length: 260Å, R_c_, width: ∼22Å) of which in particular the R_c_ did not fit with that of di-, tetra-, or hexameric coiled-coil models (R_c_: 6.2, 9.6 and 12.4 Å, respectively) (Figure 5a). Spindly^FL^ showed an overall shape and dimensions similar to Spindly^1-440^ (Figure 5a). Negative stain electron microscopy (EM) on single Spindly^1-440^ and Spindly^FL^ particles confirmed the SAXS analyses: Spindly adopts elongated shapes with some characteristic “bends” along its length (Figure 5b). The overall dimensions from EM confirmed SAXS measurements, and suggested that purified Spindly may not form a typical coiled-coil. Finally, SAXS and SEC-MALLS (Size exclusion chromatography-multi angle laser light scattering) experiments indicated that Spindly^1-440^ and Spindly^FL^ are dimeric and trimeric, respectively, in solution (Figure 5a and S5b,c).

**Figure 5:**
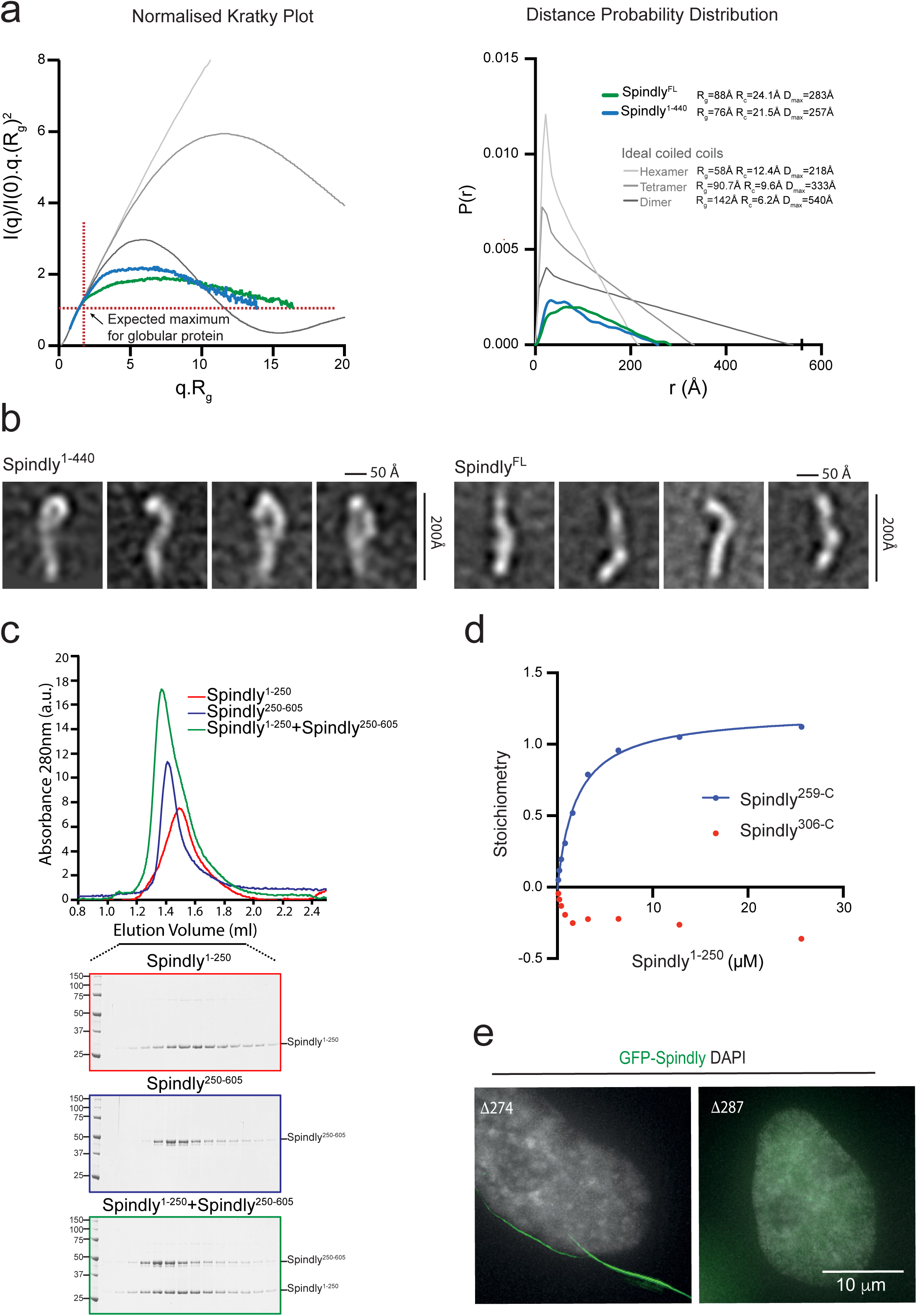
Spindly adopts a 3D configuration that impairs its ability to promote RZZS oligomerization. (a) Normalized Kratky plot (left graph) and paired distance distribution (right graph) of purified Spindly^FL^ and Spindly^1-440^. The intersection of the red dotted lines represents the peak for a globular protein irrespective of its molecular weight. Comparison with dimeric, tetrameric and hexameric coiled-coil models of similar size are shown in shades of grey. R_g_, radius of gyration; R_c_, cross-sectional radius of gyration; D_max_, maximum inter-particle distance. **(b)** Representative class averages from the negative stain EM of indicated Spindly variant proteins. **(c)** Elution profiles and SDS-PAGE of SEC experiments on Spindly^1-250^ and Spindly^250-605^ or their stoichiometric combination. AU, arbitrary units. **(d)** Surface Plasmon Resonance (SPR) analyses of the interaction between immobilized Spindly^259-C^ (blue) or Spindly^306-C^ (red) and soluble Spindly^1-250^. The response (y-axis) was normalized to the molecular weight of the analyte to yield stoichiometry of binding. **(e)** Immunofluorescence images of Spindly in interphase HeLa cells expressing the indicated truncations of GFP-Spindly.

Cross-link mass spectrometry of Spindly^FL^ supported the possibility of a more complicated Spindly fold, and revealed the existence of various long range intramolecular interactions (Figure S5c). Several of these interactions occurred around a cluster of cross-linked lysines at positions 276, 278, 283 and 284 (Figure S5c). Size exclusion chromatography showed interaction between an N-terminal fragment of Spindly (Spindly^1-250^) and a Spindly fragment encompassing C-terminal sequences (Spindly^250-605^) (Figure 5c), which was verified by Surface Plasmon resonance (SPR) (Figure 5d). This further verified intramolecular Spindly interactions and implicated the N-terminal region in these. No interaction was observed between Spindly^1-250^ and Spindly^306-605^ (Figure 5c), suggesting amino acids 259-305 have a crucial role in long-range interactions with the N-terminal Spindly helices. This was consistent with the position of the lysine cluster in the cross-link mass spectrometry analysis (Figure S5c). Notably, while a truncated Spindly lacking the N-terminal 274 amino acids (Spindly^Δ274^) retained ability to form cytoplasmic filaments in cells, this was abolished by additional removal of 13 amino acids (Spindly^Δ287^) (Figure 5e). Thus, this region that is crucial for Spindly intramolecular interactions, is also important for *in vivo* filament formation.

Together, these data support the hypothesis that the Spindly N-terminal region imposes an auto-inhibited configuration that precludes RZZ-Spindly oligomerization.

### Release of Spindly autoinhibition promotes its interaction with RZZ

We next performed SPR analyses with immobilized, purified RZZ to examine interactions of recombinant Spindly versions with the RZZ scaffold. In the absence of C-terminal farnesylation, ∼2 molecules of Spindly^FL^ weakly bound one (dimeric) molecule of RZZ with a K_D_ of ∼1 μM (Figure 6a). Farnesylation had little impact on overal interaction affinity *in vitro* but increased the number of Spindly molecules accumulating on RZZ. Similar observations were made with an alternative source of Spindly protein (Figure S6a). The farnesyl group thus appears to target Spindly to multiple sites on RZZ or to other Spindly molecules already on RZZ, under these conditions. A version of Spindly lacking the N-terminal helices (Spindly^54-605^) associated with RZZ with higher affinity (∼0.7 μM) and at higher stoichiometries: At least four molecules of Spindly could associate with RZZ. Notably, farnesylation no longer impacted interactions between RZZ and Spindly^54-605^ (Figure 6a). Removal of the N-terminal helices of Spindly therefore can achieve high occupancy and higher affinity RZZ-Spindly interactions independently of farnesylation. Based on our observation that Spindly^ΔN^ farnesylation was required for cytoplasmic filament formation (see Figure 4c) but was dispensible for Spindly^ΔN^-RZZ interactions when RZZ was on the SPR chip, we reasoned that farnesylation might facilitate initial Spindly^ΔN^- RZZ interactions that promote additional, farnesyl-independent ones. To test this, we examined if RZZ complexes concentrated on mitotic kinetochores can recruit unfarnesylated Spindly molecules. As expected, mutation of the farnesylated cysteine (C602A) or treatment with the farnesyl transferase inhibitor Lonafarnib (SCH66336) ^61^ prevented Spindly^FL^ localization and fibrous corona formation (Figure 6b-e and S6c,e). Removal of the N-terminal helices (Spindly^ΔN^) not only rescued the Spindly localization defect caused by impairing farnesylation, but also rescued fibrous corona formation (Figure 6b-d). Rescue of kinetochore localization was dependent on the 275-287 region and was independent of dynein (Figure S6c-f). We thus conclude that releasing the auto-inhibited state of Spindly, aided by C-terminal farnesylation, is necessary for formation of the fibrous corona.

**Figure 6:**
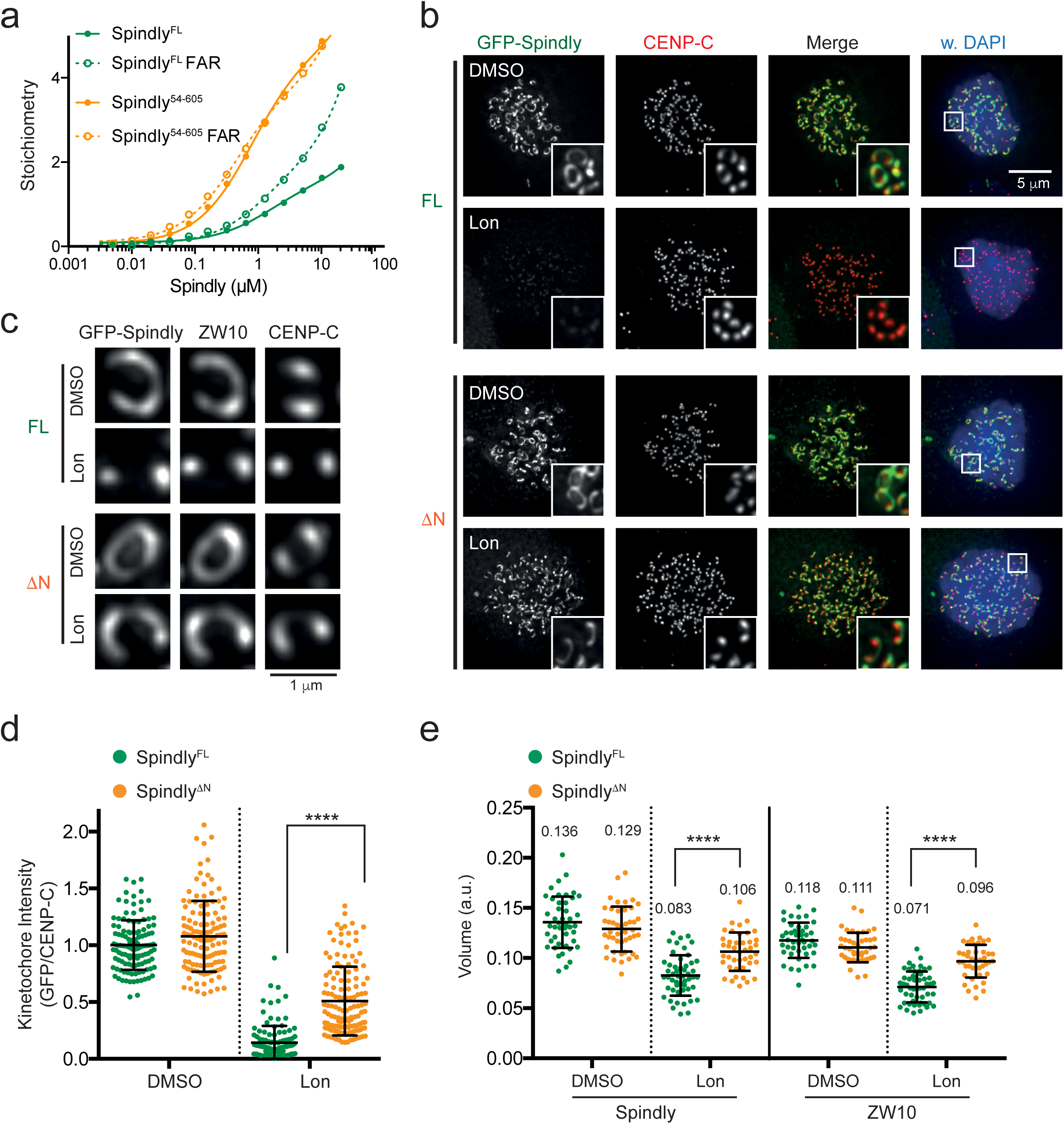
Release of Spindly autoinhibition promotes its interaction with RZZ. (a) SPR analyses of the indicated Spindly variant proteins. The response (y-axis) was normalized to the molecular weight of the analyte to yield stoichiometry of binding. **(b-e)** Immunofluorescence (b,c) and quantification of kinetochore levels (d) or volumes (e) of Spindly in HeLa expressing GFP-Spindly^FL^ or GFP-Spindly^ΔN^ and treated with nocodazole and the farnesyl transferase inhibitor Lonafarnib (Lon). The graph shows the mean fold change in kinetochore intensity (±s.d.) normalized to the values of Spindly^FL^ from three independent experiments. Each dot in D represents one cell and at least 30 cells per experiment and condition were quantified. Each dot in E represents one kinetochore and at least 10 kinetochores per experiment were quantified. Asterisks indicate significance (analysis of variance with Tukey’s multiple comparison test). ****P < 0.001.

### Activity of the kinetochore kinase MPS1 promotes fibrous corona expansion

Our data thus far suggest a mechanism for fibrous corona formation in which release of Spindly auto-inhibition enables direct interactions with RZZ and other Spindly molecules to form the corona meshwork. We deemed it likely that such a release would occur at or near kinetochores in mitosis, and reasoned that kinetochore-localized kinases are likely candidates to cause the triggering event. Whereas addition of small molecule inhibitors of Aurora B or PLK1 prior to mitotic entry caused no evident problem with fibrous corona formation in nocodazole/MG132-treated cells (Figure 7a,b), Spindly localization and the corona were substantially affected by MPS1 inhibition, to an extent similar to Spindly depletion (Figure 7b,c-f). Importantly, Spindly kinetochore levels and fibrous corona formation were rescued in MPS1-inhibited cells by deletion of the N-terminal helices of Spindly (Figure 7c-f). Thus, MPS1 is required for the release of the auto-inhibited state of Spindly. To examine if MPS1 is sufficient to trigger Spindly-dependent RZZ oligomerization, we targeted MPS1 to cytoplasmic Spindly by co-expressing GFP-Spindly and DARPin^GFP^-MPS1^Δ200^. This MPS1 variant is unable to localize to kinetochores ^62^ but can bind Spindly’s GFP tag by virtue of a GFP-binding DARPin moiety ^63^. Strikingly, when bound by active but not inactive MPS1, wild-type GFP-Spindly was able to induce interphasic filament formation (Figure 7g), just like Spindly^ΔN^ could in the absence of MPS1 activity (see Figure 4a).

**Figure 7:**
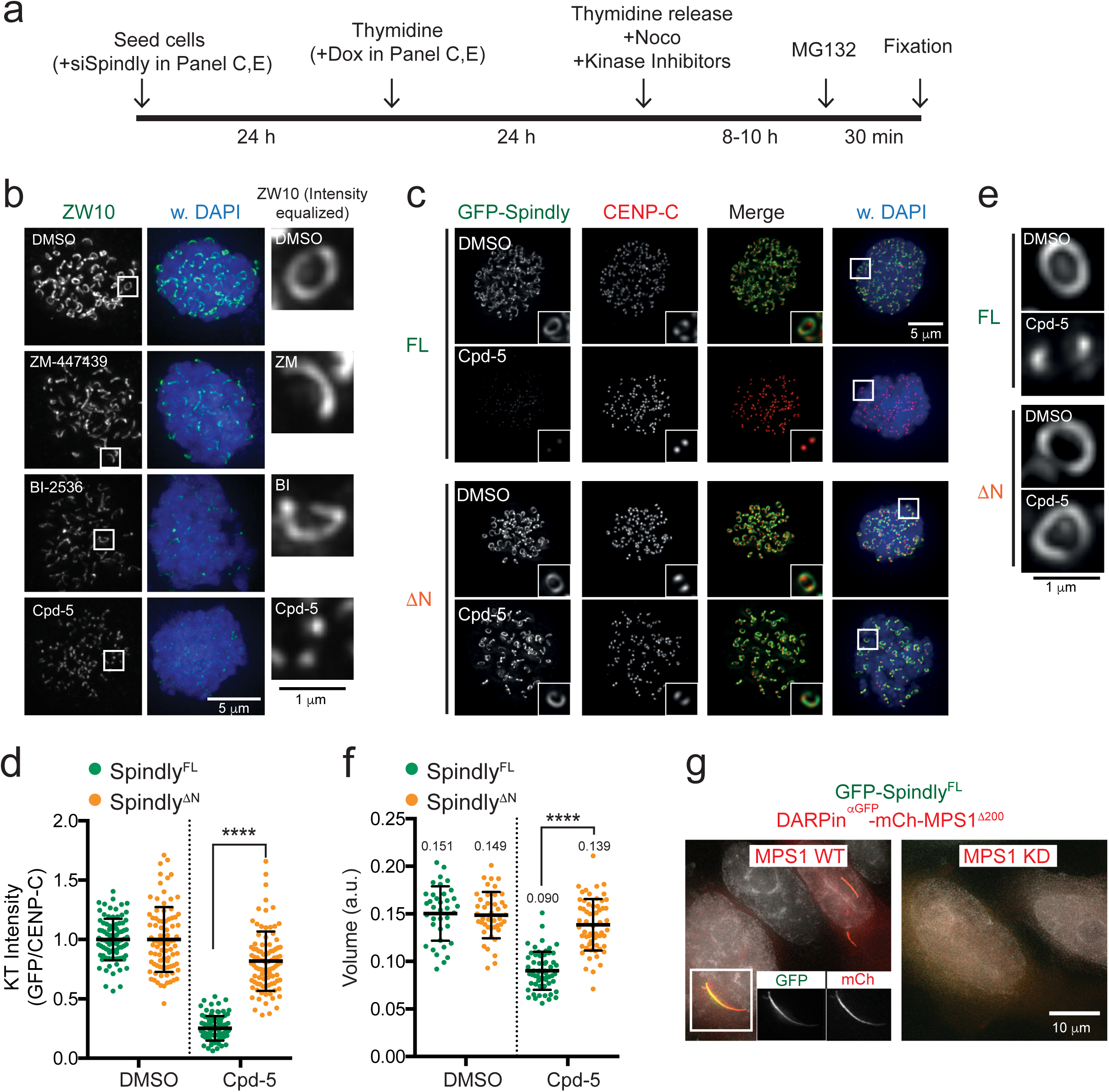
Activity of the kinetochore kinase MPS1 promotes fibrous corona expansion. (a) Timeline of the treatments with kinase inhibitors and nocodazole (Noco) of the experiments shown in b-f. **(b)** Representative images of ZW10 immunostainings of cells treated as indicated in A. The intensity levels of the zoom ins (on right) were equalized to facilitate the direct comparison of the size of the kinetochores. **(c,d)** Representative images (c) and quantification (d) of kinetochore localization of the indicated GFP-Spindly variants in HeLa cells treated with nocodazole and Cpd-5 as indicated in a. The graph shows the mean fold change in kinetochore intensity (±s.d.) normalized to the values of Spindly^FL^ from three independent experiments. Each dot represents one cell and at least 30 cells per experiment and condition were quantified. Asterisks indicate significance (analysis of variance with Tukey’s multiple comparison test). ****P < 0.001. **(e,f)** Representative images (e) and volume quantification (f) of immunostained kinetochores of HeLa cells expressing the indicated versions of GFP-Spindly and treated as indicated in a. To measure the volume, imaging acquisition was set to obtain similar mean intensity levels for the different conditions. The graph shows the mean kinetochore volume (±s.d.) of at least three independent experiments. Each dot represents one kinetochore and at least 10 kinetochores per experiment were quantified. Asterisks indicate significance (analysis of variance with Tukey’s multiple comparison test). ****P < 0.001. **(g)** Images of HeLa cells overexpressing GFP-Spindly^FL^ and an active (WT) or kinase dead (KD) version of mCherry-MPS1^Δ200^ targeted to GFP-Spindly by DARPin^GFP^.

We conclude that MPS1 kinase activity triggers the ability of Spindly to promote RZZ-Spindly oligomerization by releasing an inhibitory intra-molecular Spindly interaction.

### The fibrous corona engages in extensive lateral microtubule interactions

The persistent presence of an expanded fibrous corona in cells expressing Spindly^ΔCCS^ and Spindly^ΔN^ provided a means to examine the function of the corona and the importance of its shedding in late mitosis. Imaging of congressed chromosomes in Spindly^ΔN^-expressing cells showed that the corona of sister chromatids engaged with the sides of microtubules (Figure 8a) and had lower Astrin levels, indicative of fewer mature end-on kinetochore-microtubule interactions (Figure S7a,b) ^64^. The laterally interacting microtubules were positive for PRC1 (Figure S7c) and thus likely were anti-parallel bridging fibers (‘b’ in Figure 8a) ^65–67^. Cold treatment depolymerized PRC1 positive bundles (Figure S7c) and revealed the additional presence of stable, end-on microtubule interactions (Figure 8b and S7c). Super-resolution imaging by Expansion Microscopy (ExM) ^68^ confirmed the co-occurrence of lateral and end-on attachments on the same kinetochore (‘l’ and ‘e’ in Figure 8c) and showed an extensive fibrous corona surface capable of lateral microtubule interactions (Example 1 in Figure 8c). Live imaging of mCherry-Tubulin and GFP-Spindly^ΔN^ further showed that the corona maintained interactions with the lattices of microtubules that go through rounds of catastrophe and rescue (‘Cat’ and ‘Res’ in Figure 8d and Movies S3, S4), and with the lattices of microtubules that move away from the sister kinetochores (Figure 8d and Movies S3 and S4). The expanded fibrous corona thus efficiently captures and maintains interactions with the lattices of dynamic microtubules (Figure 8e).

**Figure 8:**
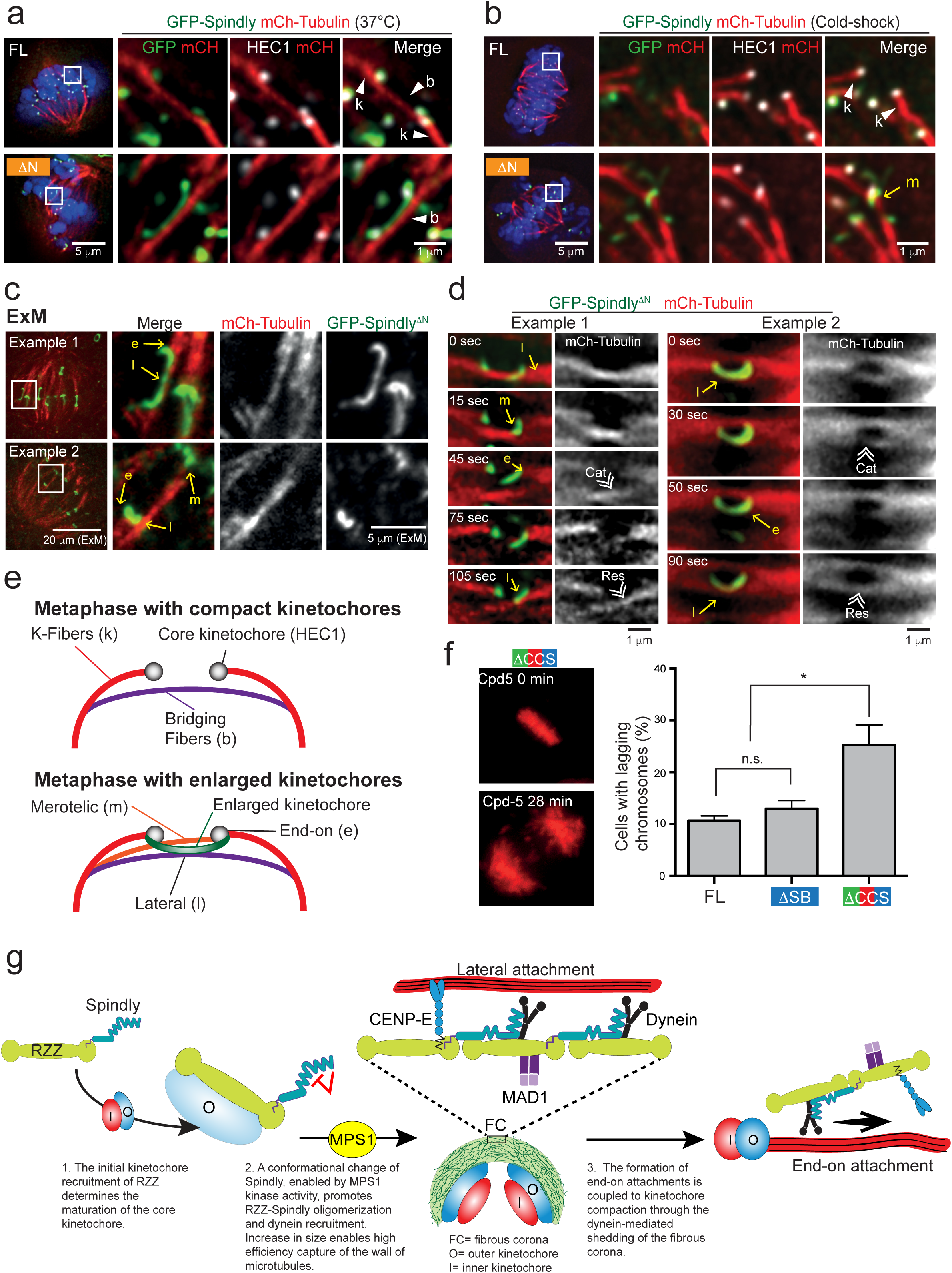
Expanded kinetochores engage in extensive lateral microtubule interactions and prevent bi-orientation. (a,b) Representative single z-plane immunofluorescence images of HeLa cells in metaphase, transfected with siRNA to Spindly and expressing the indicated GFP-Spindly variants and mCherry-tubulin. Cells were fixed at 37°C (a) or after cold-shock (b). ‘b’, bridging fibers; ‘k’, k-fibers; ‘m’ merotelic attachment. **(c)** Two examples of ExM (expansion microscopy) images of HeLa cells transfected with siRNA to Spindly and expressing GFP-Spindly^ΔN^ and mCherry-tubulin. Single z-plane images show pairs of enlarged kinetochores with simultaneous end-on (e) and lateral (l) attachments (example 1) or end-on (e), lateral (l), and merotelic (m) attachments (example 2). **(d)** Live-cell imaging of HeLa cells transfected with siRNA to Spindly and expressing GFP-Spindly^ΔN^ and mCherry-tubulin. ‘Cat’, catastrophe; ‘Res’, rescue. Maximum projections of several z-planes are shown. See also Movies S3 (Example 1) and S4 (Example2). **(e)** Cartoon summarizing the interaction of compacted or enlarged kinetochores with microtubule fibers. **(f)** Quantification of chromosome segregation errors by live-cell imaging of meptaphase HeLa cells transfected with siRNA to Spindly, expressing the indicated GFP-Spindly variants, and treated with Cpd-5. The bar graph shows the mean percentage (±s.d.) of cells showing lagging chromosomes of two independent experiments. Asterisk indicate significance (analysis of variance with Tukey’s multiple comparison test). *P < 0.01; n.s., not significant. (FL, n=222 cells; ΔSB=179 cells; ΔCCS, n=167 cells). **(g)** Model of the mechanism of kinetochore expansion and compaction performed by the axis RZZ-Spindly-dynein and MPS1. See discussion for details.

### Proper bi-orientation and chromosome segregation rely on dynamic kinetochore size regulation

We next examined the importance of fibrous corona shedding (i.e. kinetochore compaction) once microtubules are captured. We noticed that the expanded coronas of sister chromatids in Spindly^ΔN^ cells often appeared fused and frequently formed merotelic attachments (‘m’ in Figure 8b and Example 2 of Figure 8c). Live imaging captured these events: During successive cycles of capture-release, we observed short-lived events in which fibrous coronas simultaneously bound to dynamic microtubule plus ends and to stable k-fibers, briefly causing merotelic attachments (see second 15 in Example 1 of Figure 8d and Movie S3). Merotely is an important cause for chromosome segregation errors in healthy and cancerous cells ^34,69,70^. To examine if inability to shed the corona increases the frequency of chromosome segregation errors, we live imaged Spindly mutant cells undergoing anaphase, induced by treatment with an MPS1 inhibitor to bypass the SAC silencing defect resulting from absence of kinetochore dynein ^51,52^. We analyzed only cells that achieved full chromosome alignment so as not to bias for alignment problems associated with persistent coronas (Figure 8f). In contrast to cells expressing Spindly variants that allowed corona shedding (Spindly^FL^ and Spindly^ΔSB^), cells expressing Spindly^ΔCCS^ showed a high rate of lagging chromosomes in anaphase, indicative of persistent merotelic attachments.

Together these experiments show that the fibrous corona promotes lateral microtubule capture, and that subsequent corona shedding is important to establish proper, amphitelic attachments of sister chromatids.

## DISCUSSION

While the understanding of the assembly mechanisms of the core kinetochore has seen great progress in recent years ^2,71^, the nature of the fibrous corona has remained enigmatic. Here we have shown that the fibrous corona is composed of an RZZ-Spindly filamentous network. The properties of this network resemble those of other cellular polymers that assemble through a ‘collaborative’ mechanism ^72^. Collaborative filaments form on a supporting matrix, such us DNA or membranes, and frequently use hydrophobic interactions as a driving force for their assembly. By analogy, RZZ assembles on the core kinetochore in a manner dependent on BUB1 and KNL1 ^39,73^, and subsequently forms a stable filamentous structure that is maintained even when the core kinetochore is lost after inhibiting CDK1. Possibly, recruitment of BUB1 to the KMN network creates a specific physical-chemical environment that is prone to shielding by the hydrophobic fibrous corona, causing the formation of rings embracing the centromere region, much like coating proteins covering vesicles ^35^.

The RZZ-Spindly module provides the unique property of adapting kinetochore size to microtubule attachment status ^17^: In the absence of microtubules, RZZ-Spindly polymerize into a structurally stable filamentous meshwork, while microtubule binding promotes shedding of the meshwork by Spindly-dependent dynein/dynactin activity. This Spindly-centered mechanism promotes error-free chromosome birorientation by enabling efficient microtubule capture and preventing merotely by ensuring timely kinetochore compaction. Based on our observations we propose the following mechanistic model (Figure 8g): Before localizing to kinetochores, Spindly exists in an auto-inhibited state, where the N-terminal helices mask a surface required for polymerization of RZZ. Farnesylation-dependent targeting of Spindly to kinetochores by virtue of an initial interaction with ROD then enables kinetochore-localized MPS1 activity to release Spindly auto-inhibition and stimulate RZZ-Spindly polymerization ^35,43,51,54,55^. This regulation can be entirely bypassed if the N-terminal helices of Spindly are removed, suggesting that this region acts as a focal point of intra-molecular regulation. Interestingly, intramolecular interactions similar to those in Spindly have been observed for the related dynein adaptor, BICD2. In BICD2, cargo binding releases auto-inhibition, thereby allowing its N-terminal region to bind dynein-dynactin ^74^. Albeit BICD2 and Spindly share conserved short sequence motifs that affect dynein binding, our SAXS and EM data suggest an indirect role of the sequence motifs in dynein binding, consistent with a recent structural study ^75^. This raises interesting questions about how the structural organization of Spindly facilitates regulated RZZ interactions. It remains to be tested if the conformational change of Spindly is necessary to bind or activate dynein at the kinetochore. Interestingly, Spindly binds dynein efficiently but is quite heterogeneous in its ability to stimulate dynein mobility ^59^, perhaps due interference from long-range intramolecular interactions.

Several proposed events in our model of Figure 8g require further investigations. For example, the details of how MPS1 promotes release of Spindly auto-inhibition are unknown, but given that MPS1 can stimulate Spindly-RZZ filamentation in interphase it likely involves phosphorylation of one or more components of the RZZ-Spindly complex and/or one of its binding partners. Likewise, the mechanisms by which the corona binds the microtubule lattice, and the type of microtubule attachments that trigger shedding of the fibrous corona are unknown. Several microtubule-binding proteins including CENP-E, CENP-F, CLASP1, CLASP2 and dynein show localization consistent with presence at the fibrous corona ^17,19,20,27,30,76^. While dynein is important for chromosome congression ^16,77^, our data indicate that it is not essential for the interaction of kinetochores with microtubule walls. It will be important to examine which of the other microtubule-binding proteins is the primary lattice-binding activity of the fibrous corona. CENP-E is a prime candidate ^78,79^. Dynein is, on the other hand, crucial for shedding of the corona ^26,45,51,52,80^. This is likely mediated by end-on microtubule attachments, because lateral interactions are not sufficient to remove MAD1 from kinetochores ^79^. How such end-on interactions can form in the presence of the fibrous corona is unclear, but they may not need to be of the stable kind that is mediated by the NDC80 complex. We can envision [and have observed (CS and GJPLK, unpublished observations)] transient interactions between microtubule plus-ends and the fibrous corona that are sufficient to cause some corona shedding. This would not only expose NDC80 complexes, but would also remove the reported RZZ-dependent inhibition of their microtubule binding activity ^81,82^. NDC80 complexes can then engage in more stable end-on microtubule interactions that further stimulate dynein activity. Fibrous corona removal could thus promote progressive maturation of end-on attachments in a positive feedback. Such a scenario combines efficient lateral capture of microtubules with efficient transition to end-on attachments while minimizing the risk of merotelic attachments and subsequent chromosome segregation errors.

## ACKNOWLEDGMENTS

We thank members of the participating labs for suggestions and discussions. We are grateful to Andrea Murachelli for his help with critically assessing the EM data and preparing the relevant figures, to Eleonore von Castelmur, Tatjana Heidebrecht and Yoshitaka Hiruma for designing and performing initial experiments for characterising the Spindly structure, to Joshua Vaughan for help with ExM, to Reto Gassmann for sharing unpublished results and Spindly constructs, to Anko de Graaf of the Hubrecht Imaging Center, and to Iain Cheeseman, Susanne Lens and René Medema for reagents. We thank the Horizon 2020 iNEXT project (653706) for financial support and for providing access to Biophysics and EM infrastructures. This work is part of the Oncode Institute which is partly financed by the Dutch Cancer Society. Agencies that funded this work are the Netherlands Organisation for Scientific Research (NWO) (gravitation program CancerGenomiCs.nl; VICI grant (865.12.004 to GJPLK)), the Dutch Cancer Society (KWF/HUBR-11080 to GJPLK), and the European Union’s Horizon 2020 research and innovation programme under the Marie Sklodowska-Curie grant agreement No 675737 (to A.M.). V. Groenewold is supported by the Proteins@Work initiative of the Netherlands Proteomics Centre.

## Author Contributions

C.S. and G.J.P.L.K. conceived the idea for this project. C.S., G.J.P.L.K., M.A. A.P., J.K. and A.M designed the experiments and interpreted the data. C.S. performed the *in vivo* experiments. M.A. and J.K. performed the *in vitro* experiments with the help of A.F. J.F. and J.K. performed and analysed the electron microscopy experiments in cells with the help of C.S. V.G. performed the cross-linking experiments. E.T. performed the comparative sequence analysis. R.M. and J.M.C performed the electron microscopy of Spindly. C.S. and G.J.P.L.K. wrote the manuscript with the help of A.P. and A.M. and the input of the rest of authors.

## References

1. Cheeseman, I. M. The Kinetochore. Cold Spring Harb. Perspect. Biol. 6, a015826 (2014).

2. Musacchio, A. & Desai, A. A Molecular View of Kinetochore Assembly and Function. Biology (Basel). 6, 5 (2017).

3. de Wolf, B. & Kops, G. J. P. L. in Advances in Experimental Medicine and Biology 1002, 69–91 (2017).

4. van Jaarsveld, R. H. & Kops, G. J. P. L. Difference Makers: Chromosomal Instability versus Aneuploidy in Cancer. Trends in Cancer 2, 561–571 (2016).

5. Bakhoum, S. F. & Compton, D. A. Kinetochores and disease: Keeping microtubule dynamics in check! Current Opinion in Cell Biology 24, 64–70 (2012).

6. Webster, A. & Schuh, M. Mechanisms of Aneuploidy in Human Eggs. Trends in Cell Biology 27, 55–68 (2017).

7. Beach, R. R. et al. Aneuploidy Causes Non-genetic Individuality. Cell 169, 229–242.e21 (2017).

8. Santaguida, S. & Amon, A. Short- and long-term effects of chromosome missegregation and aneuploidy. Nature Reviews Molecular Cell Biology 16, 473–485 (2015).

9. Duijf, P. H. G. & Benezra, R. The cancer biology of whole-chromosome instability. Oncogene 32, 4727–4736 (2013).

10. Joglekar, A. P. & Kukreja, A. A. How Kinetochore Architecture Shapes the Mechanisms of Its Function. Current Biology 27, R816–R824 (2017).

11. Hinshaw, S. M. & Harrison, S. C. Kinetochore Function from the Bottom Up. Trends in Cell Biology 28, 22–33 (2018).

12. Etemad, B. & Kops, G. J. Attachment issues: Kinetochore transformations and spindle checkpoint silencing. Current Opinion in Cell Biology 39, 101–108 (2016).

13. Krenn, V. & Musacchio, A. The Aurora B Kinase in Chromosome Bi-Orientation and Spindle Checkpoint Signaling. Frontiers in Oncology 5, 225 (2015).

14. Sacristan, C. & Kops, G. J. P. L. P. L. Joined at the hip: Kinetochores, microtubules, and spindle assembly checkpoint signaling. Trends Cell Biol. 25, 21–28 (2015).

15. Maiato, H. The dynamic kinetochore-microtubule interface. J. Cell Sci. 117, 5461–5477 (2004).

16. Maiato, H., Gomes, A., Sousa, F. & Barisic, M. Mechanisms of Chromosome Congression during Mitosis. Biology (Basel). 6, 13 (2017). 1.

17. Magidson, V. et al. Adaptive changes in the kinetochore architecture facilitate proper spindle assembly. Nat. Cell Biol. 17, 1134–1144 (2015).

18. Wynne, D. J. & Funabiki, H. Heterogeneous architecture of vertebrate kinetochores revealed by three-dimensional superresolution fluorescence microscopy. Mol. Biol. Cell 27, 3395–3404 (2016).

19. Cooke, C. A., Schaar, B., Yen, T. J. & Earnshaw, W. C. Localization of CENP-E in the fibrous corona and outer plate of mammalian kinetochores from prometaphase through anaphase. Chromosoma 106, 446–455 (1997).

20. Yao, X., Anderson, K. L. & Cleveland, D. W. The microtubule-dependent motor centromere-associated protein E (CENP-E) is an integral component of kinetochore corona fibers that link centromeres to spindle microtubules. J. Cell Biol. 139, 435–447 (1997).

21. Thrower, D. A., Jordan, M. A. & Wilson, L. Modulation of CENP-E organization at kinetochores by spindle microtubule attachment. Cell Motil. Cytoskeleton 35, 121–133 (1996).

22. Hoffman, D. B., Pearson, C. G., Yen, T. J., Howell, B. J. & Salmon, E. D. Microtubule-dependent changes in assembly of microtubule motor proteins and mitotic spindle checkpoint proteins at PtK1 kinetochores. Mol. Biol. Cell 12, 1995–2009 (2001).

23. Magidson, V. et al. Unattached kinetochores rather than intrakinetochore tension arrest mitosis in taxol-treated cells. J. Cell Biol. 212, 307–319 (2016).

24. Wynne, D. J. & Funabiki, H. Kinetochore function is controlled by a phosphodependent coexpansion of inner and outer components. J. Cell Biol. 210, 899–916 (2015). 1.

25. Basto, R. et al. In Vivo Dynamics of the Rough Deal Checkpoint Protein during Drosophila Mitosis. Curr. Biol. 14, 56–61 (2004).

26. Griffis, E. R., Stuurman, N.& Vale, R. D. Spindly, a novel protein essential for silencing the spindle assembly checkpoint, recruits dynein to the kinetochore. J. Cell Biol. 177, 1005–1015 (2007).

27. Pereira, A. L. A. L. J. et al. Mammalian CLASP1 and CLASP2 cooperate to ensure mitotic fidelity by regulating spindle and kinetochore function. Mol. Biol. Cell 17, 4526–42 (2006).

28. Rieder, C. L. The Formation, Structure, and Composition of the Mammalian Kinetochore and Kinetochore Fiber. Int. Rev. Cytol. 79, 1–58 (1982).

29. McEwen, B. F., Hsieh, C.-E., Mattheyses, A. L. & Rieder, C. L. A new look at kineochore structure in vertebrate somatic cells using high-pressure freezing an freeze substitution. Chromosoma 107, 366–375 (1998).

30. Wordeman, L., Steuer, E. R., Sheetz, M. P. & Mitchison, T. Chemical subdomains within the kinetochore domain of isolated CHO mitotic chromosomes. J. Cell Biol. 114, 285–94 (1991).

31. Jokelainen, P. T.. The ultrastructure and spatial organization of the metaphase kinetochore in mitotic rat cells. J. Ultrastruct. Res. 19, 19–44 (1967).

32. Cassimeris, L., Rieder, C. L., Rupp, G.& Salmon, E. D. Stability of microtubule attachment to metaphase kinetochores in PtK1 cells. J. Cell Sci. 96 (Pt 1), 9–15 (1990).

33. McEwen, B. F., Arena, J. T., Frank, J.& Rieder, C. L. Structure of the Colcemid-treated PtK1 kinetochore outer plate as determined by high voltage electron microscopic tomography. J. Cell Biol. 120, 301–312 (1993). 1.

34. Lampson, M. & Grishchuk, E. Mechanisms to Avoid and Correct Erroneous Kinetochore-Microtubule Attachments. Biology (Basel). 6, 1 (2017).

35. Mosalaganti, S. et al. Structure of the RZZ complex and molecular basis of its interaction with Spindly. J. Cell Biol. 216, 961–981 (2017).

36. Karess, R. Rod-Zw10-Zwilch: A key player in the spindle checkpoint. Trends Cell Biol. 15, 386–392 (2005).

37. Kops, G. J. P. L. et al. ZW10 links mitotic checkpoint signaling to the structural kinetochore. J. Cell Biol. 169, 49–60 (2005).

38. Buffin, E., Lefebvre, C., Huang, J., Gagou, M. E. & Karess, R. E. Recruitment of Mad2 to the kinetochore requires the Rod/Zw10 complex. Curr. Biol. 15, 856–861 (2005).

39. Caldas, G. V. et al. The RZZ complex requires the N-terminus of KNL1 to mediate optimal Mad1 kinetochore localization in human cells. Open Biol. 5, 150160 (2015).

40. Silió, V., McAinsh, A. D. & Millar, J. B. KNL1-Bubs and RZZ Provide Two Separable Pathways for Checkpoint Activation at Human Kinetochores. Dev. Cell 35, 600–613 (2015).

41. Défachelles, L. et al. RZZ and Mad1 dynamics in Drosophila mitosis. Chromosom. Res. 23, 333–342 (2015).

42. Starr, D. A. et al. Conservation of the centromere/kinetochore protein ZW10. J. Cell Biol. 138, 1289–1301 (1997).

43. Gama, J. B. et al. Molecular mechanism of dynein recruitment to kinetochores by the Rod-Zw10-Zwilch complex and Spindly. J. Cell Biol. 216, 943–960 (2017). 1.

44. Çivril, F. et al. Structural analysis of the RZZ complex reveals common ancestry with multisubunit vesicle tethering machinery. Structure 18, 616–626 (2010).

45. Howell, B. J. et al. Cytoplasmic dynein/dynactin drives kinetochore protein transport to the spindle poles and has a role in mitotic spindle checkpoint inactivation. J. Cell Biol. 155, 1159–1172 (2001).

46. Wojcik, E. et al. Kinetochore dynein: Its dynamics and role in the transport of the Rough deal checkpoint protein. Nat. Cell Biol. 3, 1001–1007 (2001).

47. Famulski, J. K., Vos, L. J., Rattner, J. B. & Chan, G. K. Dynein/dynactin-mediated transport of kinetochore components off kinetochores and onto spindle poles induced by Nordihydroguaiaretic acid. PLoS One 6, e16494 (2011).

48. Mahale, S. P., Sharma, A.& Mylavarapu, S. V. S. Dynein light intermediate chain 2 facilitates the metaphase to anaphase transition by inactivating the spindle assembly checkpoint. PLoS One 11, e0159646 (2016).

49. Sivaram, M. V. S., Wadzinski, T. L., Redick, S. D., Manna, T.& Doxsey, S. J. Dynein light intermediate chain 1 is required for progress through the spindle assembly checkpoint. EMBO J. 28, 902–914 (2009).

50. Mische, S. et al. Dynein light intermediate chain: an essential subunit that contributes to spindle checkpoint inactivation. Mol. Biol. Cell 19, 4918–29 (2008).

51. Barisic, M. et al. Spindly/CCDC99 is required for efficient chromosome congression and mitotic checkpoint regulation. Mol. Biol. Cell 21, 1968–1981 (2010).

52. Gassmann, R. et al. Removal of Spindly from microtubule-attached kinetochores controls spindle checkpoint silencing in human cells. Genes Dev. 24, 957–971 (2010).

53. Ying, W. C. et al. Mitotic control of kinetochore-associated dynein and spindle orientation by human Spindly. J. Cell Biol. 185, 859–874 (2009).

54. Moudgil, D. K. et al. A novel role of farnesylation in targeting a mitotic checkpoint protein, human spindly, to kinetochores. J. Cell Biol. 208, 881–896 (2015).

55. Holland, A. J. et al. Preventing farnesylation of the dynein adaptor Spindly contributes to the mitotic defects caused by farnesyltransferase inhibitors. Mol. Biol. Cell 26, 1845–1856 (2015).

56. Tromer, E., Bade, D., Snel, B.& Kops, G. J. P. L. Phylogenomics-guided discovery of a novel conserved cassette of short linear motifs in BubR1 essential for the spindle checkpoint. Open Biol. 6, 1–11 (2016).

57. van Hooff, J. J., Tromer, E., van Wijk, L. M., Snel, B.& Kops, G. J. Evolutionary dynamics of the kinetochore network in eukaryotes as revealed by comparative genomics. EMBO Rep. 18, 1559–1571 (2017).

58. Gassmann, R. et al. A new mechanism controlling kinetochore-microtubule interactions revealed by comparison of two dynein-targeting components: SPDL-1 and the Rod/Zwilch/Zw10 complex. Genes Dev. 22, 2385–2399 (2008).

59. McKenney, R. J., Huynh, W., Tanenbaum, M. E., Bhabha, G. & Vale, R. D. Activation of cytoplasmic dynein motility by dynactin-cargo adapter complexes. Science 345, 337–341 (2014).

60. Starr, D. A., Williams, B. C., Hays, T. S. & Goldberg, M. L. ZW10 Helps Recruit Dynactin and Dynein to the Kinetochore. J. Cell Biol. 142, 763–774 (1998).

61. Njoroge, F. G. et al. (+)-4-[2-[4-(8-Chloro-3,10-dibromo-6,11-dihydro-5H-1. benzo[5,6]cyclohepta[1,2-b]-pyridin-11(R)-yl)-1-piperidinyl]-2-oxo-ethyl]-1-piperidinecarboxamide (SCH-66336): A Very Potent Farnesyl Protein Transferase Inhibitor as a Novel Antitumor Agent. J. Med. Chem. 41, 4890–4902 (1998).

62. Nijenhuis, W. et al. A TPR domain-containing N-terminal module of MPS1 is required for its kinetochore localization by Aurora B. J. Cell Biol. 201, 217–231 (2013).

63. Brauchle, M. et al. Protein interference applications in cellular and developmental biology using DARPins that recognize GFP and mCherry. Biol. Open 3, 1252–1261 (2014).

64. Shrestha, R. L. & Draviam, V. M. Lateral to end-on conversion of chromosome-microtubule attachment requires kinesins cenp-e and MCAK. Curr. Biol. 23, 1514–1526 (2013).

65. Polak, B., Risteski, P., Lesjak, S. & Tolić, I. M. PRC1-labeled microtubule bundles and kinetochore pairs show one-to-one association in metaphase. EMBO Rep. 18, 217–230 (2017).

66. Kajtez, J. et al. Overlap microtubules link sister k-fibres and balance the forces on bi-oriented kinetochores. Nat. Commun. 7, 10298 (2016).

67. Simunić, J. & Tolić, I. M. Mitotic Spindle Assembly: Building the Bridge between Sister K-Fibers. Trends in Biochemical Sciences 41, 824–833 (2016).

68. Chozinski, T. J. et al. Expansion microscopy with conventional antibodies and fluorescent proteins. Nat. Methods 13, 485–488 (2016).

69. Gregan, J., Polakova, S., Zhang, L., Tolić-Nørrelykke, I. M. & Cimini, D. Merotelic kinetochore attachment: Causes and effects. Trends in Cell Biology 21, 374–381 (2011).

70. Cimini, D. et al. Merotelic kinetochore orientation is a major mechanism of aneuploidy in mitotic mammalian tissue cells. J. Cell Biol. 152, 517–527 (2001).

71. Pesenti, M. E., Weir, J. R. & Musacchio, A. Progress in the structural and functional characterization of kinetochores. Current Opinion in Structural Biology 37, 152–163 (2016).

72. Ghosal, D. et al. Collaborative protein filaments. EMBO J. 34, 2312–2320 (2015).

73. Zhang, G., Lischetti, T., Hayward, D. G. & Nilsson, J. Distinct domains in Bub1 localize RZZ and BubR1 to kinetochores to regulate the checkpoint. Nat. Commun. 6, 7162 (2015).

74. Hoogenraad, C. C. & Akhmanova, A. Bicaudal D Family of Motor Adaptors: Linking Dynein Motility to Cargo Binding. Trends Cell Biol. 26, 327–340 (2016).

75. Urnavicius, L. et al. Cryo-EM shows how dynactin recruits two dyneins for faster movement. Nature 554, 202–206 (2018).

76. Maiato, H. et al. Human CLASP1 is an outer kinetochore component that regulates spindle microtubule dynamics. Cell 113, 891–904 (2003).

77. Barisic, M., Aguiar, P., Geley, S. & Maiato, H. Kinetochore motors drive congression of peripheral polar chromosomes by overcoming random arm-ejection forces. Nat. Cell Biol. 16, 1249–1256 (2014).

78. Kapoor, T. M. et al. Chromosomes can congress to the metaphase plate before biorientation. Science 311, 388–91 (2006).

79. Kuhn, J. & Dumont, S. Spindle assembly checkpoint satisfaction occurs via end-on but not lateral attachments under tension. J. Cell Biol. 216, 1533–1542 1. (2017).

80. Varma, D., Monzo, P., Stehman, S. A. & Vallee, R. B. Direct role of dynein motor in stable kinetochore-microtubule attachment, orientation, and alignment. J. Cell Biol. 182, 1045–1054 (2008).

81. Cheerambathur, D. K., Gassmann, R., Cook, B., Oegema, K. & Desai, A. Crosstalk Between Microtubule Attachment Complexes Ensures Accurate Chromosome Segregation. Science 342, 1239–1242 (2013).

82. Amin, M. A., Mckenney, R. J. & Varma, D. Antagonism between the dynein and Ndc80 complexes at kinetochores controls the stability of kinetochore-microtubule attachments during mitosis. J. Biol. Chem. jbc.RA117.001699 (2018). doi:10.1074/jbc.RA117.001699

